# Binding Paths: Describing Small Molecule Interactions with Disordered Proteins via a Markov State Model

**DOI:** 10.1101/2025.11.19.688850

**Authors:** Adelie Louet, Gerhard Hummer, Michele Vendruscolo

## Abstract

Disordered proteins are challenging targets for drug discovery because they lack well-defined binding pockets. Although small molecules can form relatively stable complexes with disordered proteins, the highly dynamic nature of these proteins complicates the understanding of their binding mechanism. To address this problem, we analyze the binding of the small molecule 10074-G5 to Aβ42, which results in the formation of a disordered complex. We describe the binding mechanism in terms of binding paths, which are stochastic trajectories along which small molecule diffuse across disordered protein surfaces, forming transient contacts with overlapping groups of residues. To identify these binding paths, we define them as realizations of a stochastic process defined by a Markov State Model (MSM). The MSM is built from distinct states of the disordered complex and their corresponding transition probabilities, enabling both the dynamic mapping of binding hotspots and targetable regions where static pockets cannot be defined-extending the notion of binding pockets to disordered systems - and the quantitative calculation of binding affinities. The visualization of the MSM via knowledge graphs provides an intuitive representation of the binding paths. We further validated this approach across four additional systems, comprising C-terminal α-synuclein with three distinct small-molecule binders, and the full-length peptide (140 residues) with binder fasudil. By generalizing the concept of static binding pockets to dynamic binding paths, our approach rationalizes small-molecule recognition by disordered proteins and establishes a framework for identifying druggable regions on systems that were previously considered intractable, providing insights for future drug design programs.

## Introduction

Disordered proteins make up about one third of the human proteome and play essential roles in signaling, regulation, and molecular recognition^1-4^. Although their dysregulation is central to a wide range of human diseases, despite recent progress they are still considered undruggable - a label that reflects our current lack of concepts and methods to discover effective drugs targeting them^5-16^.

Over the past two decades, a wide range of computational structure-based drug-discovery pipelines, including virtual screening, molecular docking, molecular-dynamics refinement and free-energy perturbation, have flourished because many disease-relevant proteins present well-defined binding pockets^17-22^. In this type of approach, a three-dimensional pocket is first identified in an experimentally-determined structure of a target protein. Then, chemical libraries typically containing millions of compounds are screened in silico, scored for shape and electrostatic complementarity, and the top-ranked hits are iteratively optimised with physics-based free-energy calculations before experimental validation. This strategy has generated approved drugs, from kinase inhibitors in oncology to the pharmacological chaperone tafamidis, which fits into a thyroid-hormone-like cleft in transthyretin and prevents its misfolding^23,24^.

In the case of disordered proteins, although stable binding pockets do not typically exist^5-15^, it has been recently shown that the binding of small molecules can arise through the formation of ordered tertiary contacts upon binding^25,26^. Using molecular dynamics simulations validated by nuclear magnetic resonance (NMR) spectroscopy, small molecules can be identified targeting the disordered N-terminal transactivation domain of the androgen receptor, and shown to induce the formation of compact, *α*-helical, molten-globule-like conformations ^25,26^.

Advances have been made also in cases where the bound state remains disordered. NMR relaxation dispersion and small-angle X-ray scattering were used to show that ligand binding to the disordered D2 domain of p27^Kip1^ causes a population shift among pre-existing conformations within a disordered structural ensemble^27^. These findings support a model in which binding arises from redistribution of populations across transient, low-energy states already present in the unbound ensemble. This behavior contrasts with models that emphasize ligand-induced disorder expansion^6^, and it highlights the dynamic complexity of ligand recognition for disordered protein.

Additional insight about the dynamic nature of small-molecule interactions with disordered proteins comes from recent long-timescale molecular dynamics simulations and NMR studies of fasudil binding to α-synuclein^28^. Fasudil was shown to preferentially interact with the disordered C-terminal region of α-synuclein through a network of weak, transient charge-charge and π-stacking interactions^28^. Rather than forming a single, stable binding mode, fasudil exhibited a dynamic shuttling mechanism, transitioning among multiple complementary interactions without locking into a fixed pose. This dynamic structural ensemble was consistent with residue-specific NMR chemical shift perturbations and validated through the design of fasudil analogs spanning different binding affinities^28^.

Further experimental evidence substantiates the notion that small molecules can interact with disordered proteins in highly dynamic ways. Using ^19^F NMR spin relaxation techniques, it was shown that 5-fluoroindole binds to disordered domains of the hepatitis C virus protein NS5A with *μ*M affinity, yet remains mobile in its bound state, with a rotational correlation time of just 46 ps, which is only marginally longer than in its free form^29^. These findings provide an experimental confirmation of a bound state characterized by persistent, rapid fluctuations, supporting the emerging view that small molecule interactions with disordered proteins are not governed by single, stable pockets but rather by ensembles of transient, loosely defined contacts^6,28-30^.

A promising example is provided by the interaction between the monomeric state of Aβ42 and the small-molecule inhibitor 10074-G5 (referred to as G5 in the following)^31,32^. It was shown G5 binds the monomeric state of Aβ42 with low *μ*M affinity, thereby inhibiting its aggregation^31,32^. It was suggested that the decrease in free energy upon formation of the complex between Aβ42 and G5 is favoured by an increase in the conformational entropy of Aβ42, which in turn increases the level of disorder of Aβ42 in the bound state^31,32^. At the same time, G5 was found to form transient interactions, including hydrophobic contacts, charge-charge interactions, aromatic π-π stacking, and hydrogen bonding^31,32^. This system was selected for modeling in our pipeline to identify targetable regions in disordered proteins.

Our results support a binding mechanism in which a small molecule diffuses along the surface of the protein by binding a series of transiently formed binding pockets. This disordered binding mechanism lends itself naturally to a description in terms of a Markov state model (MSM), where the states represent the transient binding pockets, and the transition probabilities correspond to the rate constants for ligand movement between these pockets. To implement an MSM, we integrated molecular dynamics simulations, stochastic analysis of residue contacts, and network-based modeling. This approach resulted in the characterisation of binding paths, which are the stochastic routes that a small molecule traces across a disordered surface, as it sequentially engages overlapping clusters of residues forming transient contacts. Importantly, this framework enables the identification of dynamic binding hotspots and targetable regions on intrinsically disordered proteins—regions that cannot be captured by conventional static pocket-based descriptions. Our results indicate that the conceptual shift from static binding pockets to dynamic binding paths enables an accurate understanding of how small molecules interact with disordered proteins.

## Results

### VAMPNet clustering of binding pockets of A*β*42

To explore whether discrete binding pockets could be identified in the interaction between Aβ42 and G5, we initially applied a deep learning-based clustering approach using VAMPNet^33^. This method is designed to learn optimal representations of the slow dynamical modes of a system by training a neural network on time-lagged molecular dynamics data, thereby enabling the identification of metastable states suitable for constructing an MSM.

We investigated whether VAMPNet could be used to cluster the ligand-protein bound configurations by providing as input the time series of distances between the center of mass of the ligand and the Cα atoms of Aβ42. A two-level hierarchical clustering procedure was employed, with each level producing four clusters, ultimately yielding a total of sixteen discrete states. Despite this effort, the clustering results revealed a significant limitation: over 90% of the frames were assigned to a single dominant state (state 16) (**Figures 1** and **S2**). This outcome suggested that the application of VAMPNet to the specific problem that we studied was not readily capable of distinguishing distinct ligand binding configurations within the conformational ensemble of Aβ42. The result may reflect either a limited sensitivity of the method to subtle variations in contact patterns, or an absence of well-defined and kinetically separable binding pockets in such a highly dynamic system.

**Figure 1.**
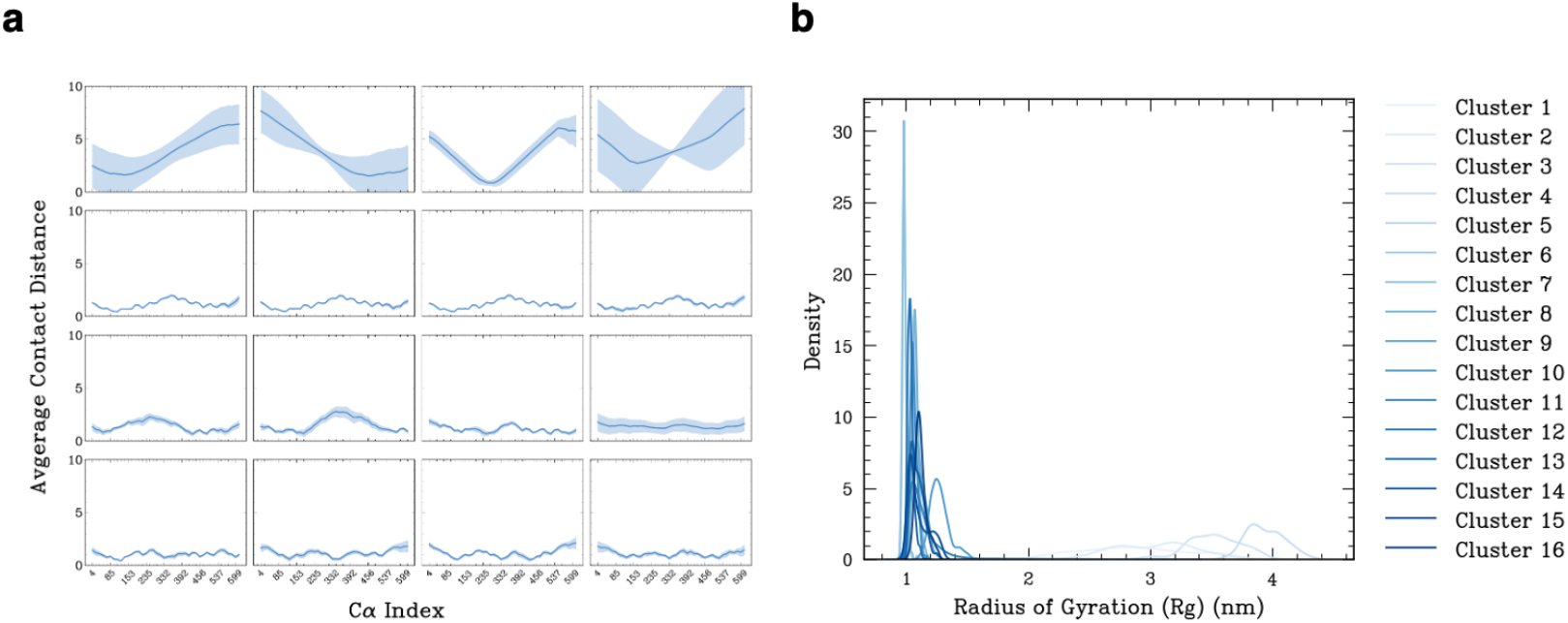
VAMPNet clustering of the binding pockets of A*β*42. **(a)** The average distance between the ligand and each Cα atom of Aβ42 shows varying levels of correlation across clusters. Smaller clusters exhibit stronger correlations, while the dominant cluster (primarily Cluster 16) shows weaker correlations and higher standard deviation from the mean. **(b)** Radius of gyration values also display substantial variability among the larger clusters, reflecting their greater structural heterogeneity.

### Stochastic nature of the binding between G5 and A*β*42

To better understand the challenges in clustering binding pockets in a disordered protein, we analyzed how G5 diffuses along the surface of Aβ42 by forming transient contacts with different side chains. This behavior was characterized by generating a time series that tracks whether each individual residue is in contact with G5 at every frame of a molecular dynamics trajectory (**Figure 2**). The resulting time series revealed a stochastic pattern of interactions, consistent with previous reports that G5 does not preferentially bind to a specific, well-defined site, such as a stable cavity or cleft, typically associated with structured proteins ^31,32^. Instead, the ligand dynamically diffuses across the Aβ42 surface, displaying generalized affinity for multiple residues rather than settling into a single, persistent binding pocket. This behavior highlights the limitations of traditional clustering approaches, which assume discrete, metastable binding states, and underscores the need for models that can account for the continuous and probabilistic nature of binding to disordered proteins.

**Figure 2.**
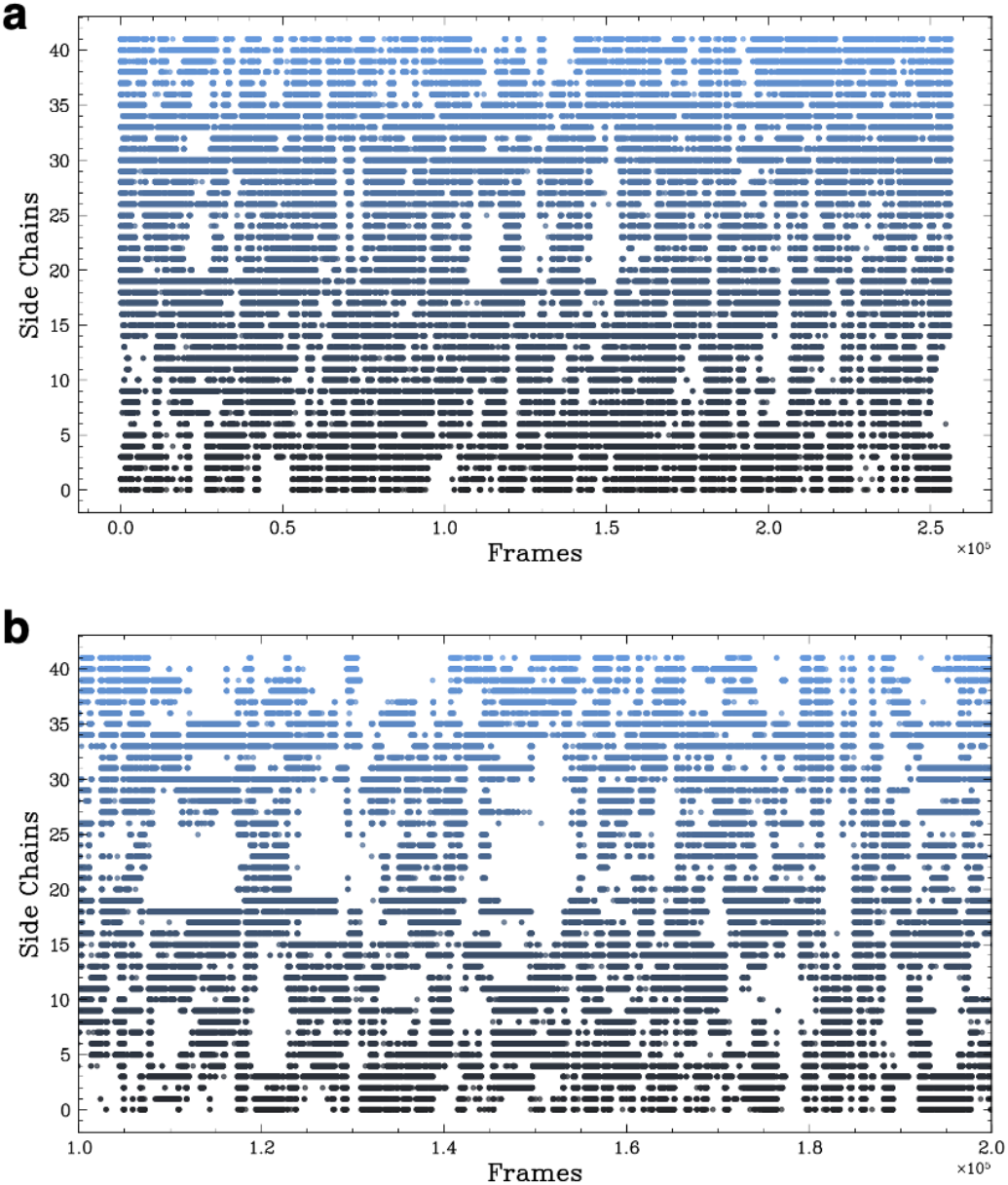
Time series of the binding of G5 to A*β*42. **(Left)** The x-axis shows the simulation time steps, and the y-axis indicates whether G5 is in contact with each residue of Aβ42. **(Right)** A subset of the trajectory (frames 1,000,000 to 2,000,000) highlights the frequent overlap and crossover between multiple residues contacting the ligand simultaneously, illustrating the dynamic and non-specific nature of binding.

### Residue-level mapping of the stochastic binding using a handoff matrix

From the analysis above, two key observations emerged. First, the inability to create discrete clusters appears to arise from the vast number of states in which disordered proteins exist. Second, a dynamic shuttling mechanism suggests that ligands diffuse rapidly on the protein surface without populating stable binding pockets. Our approach involved exploring three-dimensional and highly dynamic methodologies, focusing on the continuous interaction over the trajectory rather than isolated snapshots at individual time points.

To better understand how a small molecule ligand dynamically interacts with a disordered protein, we developed a representation that captures how the ligand transitions between residue contacts over time. We refer to this representation as the handoff matrix, which encodes the frequency with which the ligand moves from one residue to another during the molecular dynamics simulations. Specifically, for Aβ42 the handoff matrix is a 42×42 matrix *H*, where each element *H*_*ij*_ represents the number of observed transitions in which the ligand is in contact with residue *i* at one time step and with residue *j* at the subsequent time step **(Figure 3)**. This representation enabled us to capture the inherently stochastic behavior of G5 as it diffuses across the disordered surface of Aβ42, forming and breaking transient contacts with different residues. Importantly, this representation shifts the perspective from discrete, static binding events to a dynamic process in which interactions are continuously redistributed across the protein.

**Figure 3.**
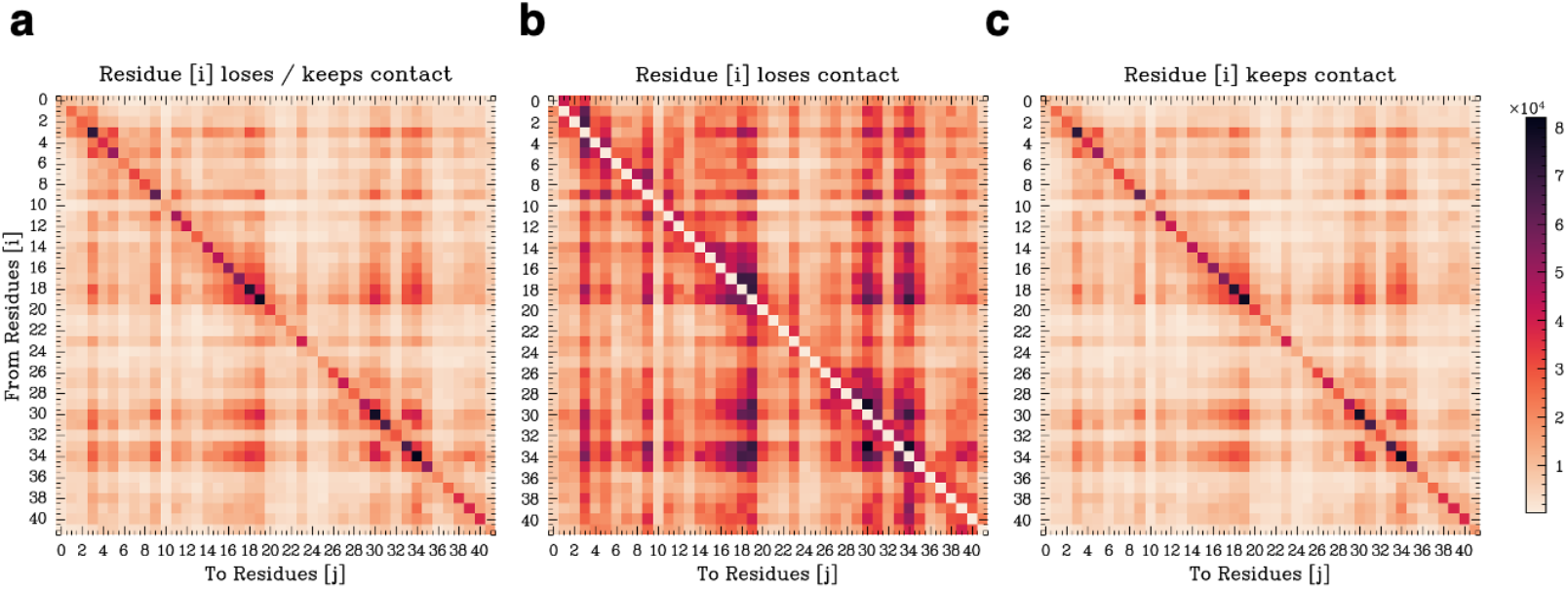
Handoff matrices for the binding of G5 to A*β*42. **(a)** Case A: A handoff occurs from residue *i* to residue *j*, regardless of whether residue *i* maintains its interaction with the ligand. **(b)** Case B: Residue *i* loses contact with the ligand while residue *j* gains contact, representing a true handoff. **(c)** Case C: Residue *i* retains its interaction, and residue *j* forms a new contact, indicating an additional interaction rather than a replacement.

To characterize the nature of these transitions, we analyzed the handoff events in terms of three specific cases. In case A, we recorded all transitions in which residue i was in contact with the ligand at time *t*, and residue *j* was in contact at time *t+1*, regardless of whether residue *i* retained its contact **(Figure 3A)**. Case B restricted this calculation to events in which residue *i* lost its contact while residue *j* gained a contact, representing a strict handoff **(Figure 3B)**. Case C captured scenarios in which residue *i* maintained its contact while residue *j* was newly engaged, highlighting additive interactions rather than replacements **(Figure 3C)**.

The resulting handoff matrix reveals key binding hotspots on the Aβ42 surface. These are regions where the ligand frequently transitions between specific pairs of residues. These hotspots reflect areas of elevated interaction probability and serve as a dynamic analogue to classical binding pockets. By averaging over matrix elements corresponding to specific groups of residues, we can define effective transition probabilities that are then incorporated into the MSM framework. This approach thus provides a mechanistically grounded basis for identifying targetable regions on disordered proteins and for describing the binding behavior of a ligand in terms of continuous surface diffusion rather than discrete docking.

### Definition of a Markov state model of the stochastic binding

To capture the dynamics of ligand binding to Aβ42, we constructed an MSM based on the transitions between residue clusters that interact with the ligand over time. In contrast to traditional approaches that define states based on global protein conformations or static binding pockets, we defined states as n-plets, which are sets of residues simultaneously in contact with the ligand at a given time point. We systematically enumerated all residue combinations of size two to five (i.e., doublets to quintuplets) that formed simultaneous contacts with the ligand throughout the simulation. Each unique combination was treated as a distinct MSM state. As the simulation progressed, we tracked how the ligand transitioned between these states by recording which sets of residues lost contact and which new sets gained contact at each time step **(Figure 4A)**.

**Figure 4.**
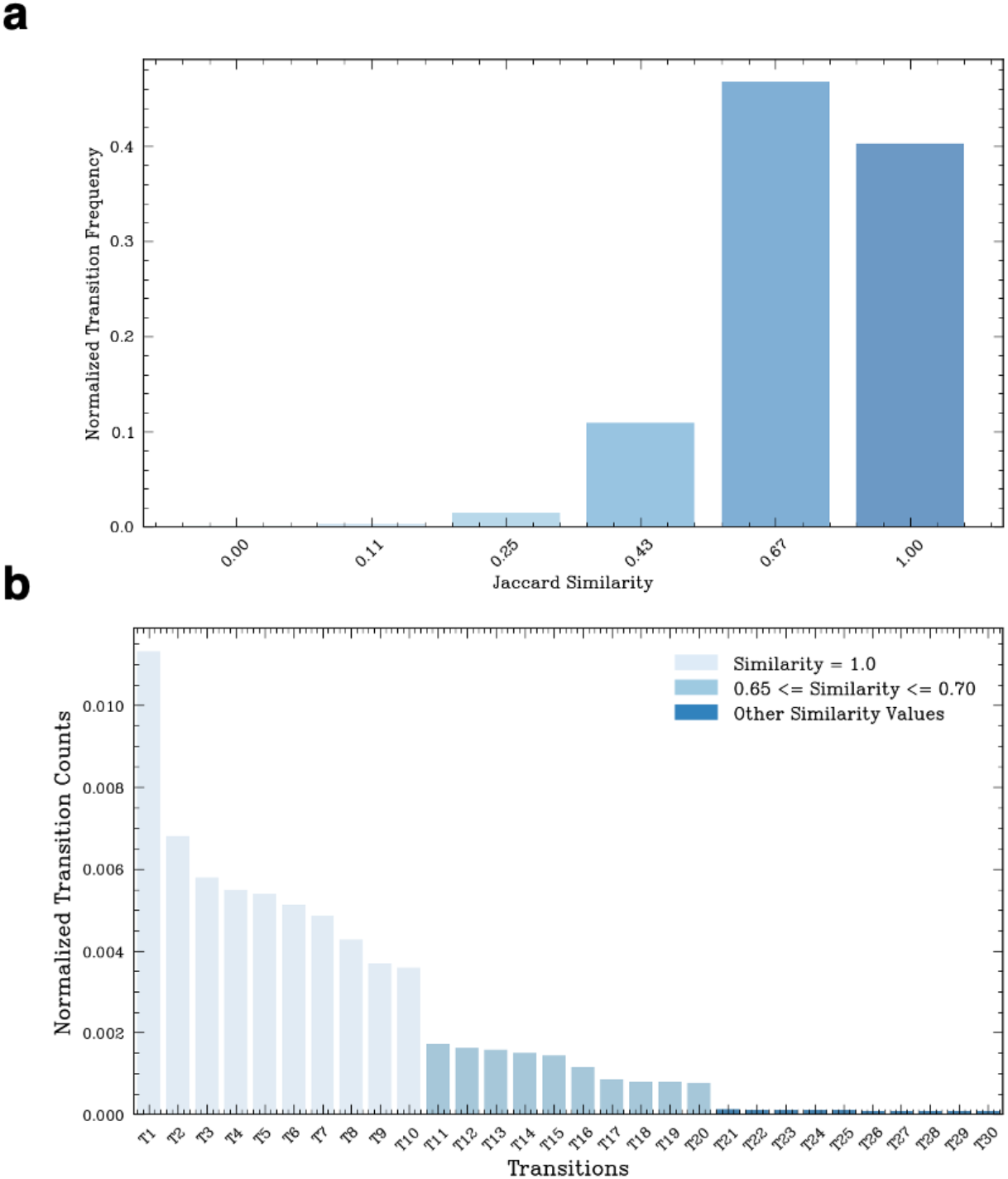
Definition of the n-plets and of the transitions between them. **(a)** Distribution of transition frequencies between quintuplets, categorized by their pairwise similarity scores. This analysis distinguishes between high-similarity glides and low-similarity jumps, revealing that transitions between dissimilar states are rare. **(b)** Transitions between quintuplets with less than 65% similarity occur with very low frequency, supporting a gliding mechanism of ligand movement on the protein surface rather than abrupt hopping between distant regions (labels corresponding to quintuplet states in **Table S2**).

This approach allowed us to construct a transition matrix, where each element represents the frequency of transitions between specific n-plets. The model distinguishes between two types of transitions: gliding transitions, where the ligand moves between similar clusters (e.g., with high residue overlap), and hopping transitions, where the ligand jumps between dissimilar regions on the protein surface. To quantify this distinction, we computed the Jaccard similarity between consecutive n-plets. We found that most transitions occurred between highly similar residue sets (Jaccard similarity ≥ 0.66), supporting a gliding mechanism of ligand diffusion across the protein **(Figure 4B)**. In contrast, transitions between dissimilar n-plets were rare (approximately 7%), indicating that long-range hops are infrequent **(Figure 4B)**.

By modeling the stochastic movement of the ligand between overlapping contact clusters, this MSM framework provides a state-based representation of the binding process, well suited to the disordered and dynamic nature of Aβ42. The resulting model offers kinetic and thermodynamic insights into how a small molecule engages with the disordered protein surface.

### Master equation analysis of the Markov state model

To quantitatively describe the binding dynamics captured by the MSM, we analyzed the system using a master equation formalism. This framework enabled us to compute how the probability distribution of ligand-bound states evolves over time, based on the transition rates between states. Each MSM state corresponds to an n-plet (for example a quintuplet) of residues simultaneously in contact with the ligand. The full set of these states defines the space in which the ligand diffuses. For Aβ42, when considering quintuplets, this procedure identified a total of 13,034 states. The transitions between these states, recorded from a molecular dynamics trajectory, define the transition rate matrix used in the master equation. The master equation describes the time evolution of the probabilities of the states of the systems

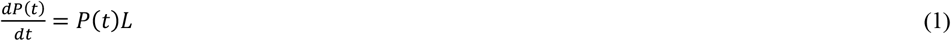

where P(t) = [*p*_*i*_(t), …, *p*_*n*_(t)] is the row vector of state probabilities *p*_*i*_(t), at time *t*, and *L* is the generator matrix, whose off-diagonal entries encode the transition rates between states and whose rows sum to zero. The long-time behavior of the system converges to a stationary distribution *P*_*eq*_ = [*p*_1,*eq*_, …, *p*_*n,eq*_], which corresponds to the equilibrium population of each state. We note that although molecular dynamics simulations generate data at discrete time intervals, we employ a continuous-time MSM to describe the binding dynamics. Starting from a discrete molecular dynamics trajectory, we first identify transitions between contact-based states separated by a chosen lag time. These transitions are used to construct a discrete transition probability matrix *T*, which captures the stochastic evolution of the system over that lag time. To obtain a time-continuous description of the dynamics, we then estimate the generator matrix *L* of the corresponding continuous-time MSM (see Eq. 3 in Methods). This approach allowed us to model the time evolution of state populations via a master equation, providing access to both kinetic and thermodynamic properties of the system. A continuous formulation of this type is particularly advantageous for capturing the inherently dynamic nature of disordered protein-ligand interactions, as it avoids artifacts associated with a fixed time resolution and enables interpolation across arbitrary time scales.

To validate the accuracy of the MSM, we performed a Chapman-Kolmogorov (CK) test, comparing the predicted evolution of state probabilities with the actual data from the simulation across multiple lag times. We found that the agreement between predicted and observed dynamics improved as the size of the residue clusters increased, with quintuplets providing the most reliable model **(Figure 5)**. This result confirms that larger n-plets more accurately capture the spatial and temporal resolution of the binding process.

**Figure 5.**
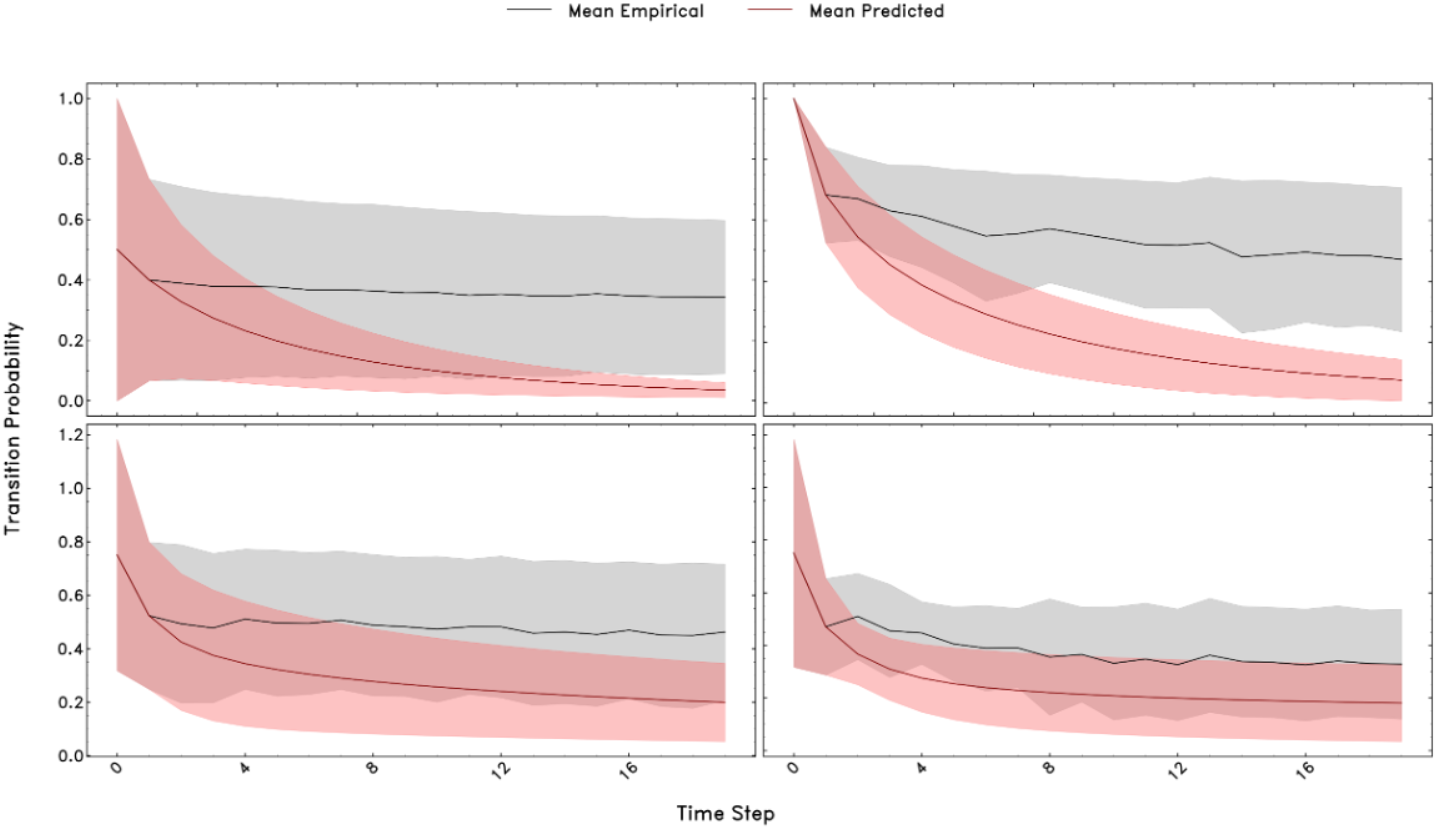
Chapman-test of the MSM. Agreement between predicted and empirical transition probabilities improves with increasing n-plet size (upper left: 2-plet, upper right: 3-plet, lower left: 4-plet, lower right: 5-plet). Larger residue clusters (e.g., quintuplets) exhibit better consistency with the Markovian assumption, supporting their use in constructing stable MSMs.

### Visualization of the MSM through knowledge graphs

To visualize the MSM, we constructed a knowledge graph, which is a type of network diagram that represents entities (nodes) and their relationships (edges). In our context, each node corresponds to a quintuplet of residues simultaneously in contact with the ligand, representing a unique state in the Markov model (**Figure 6A**). Edges between nodes indicate observed transitions between these states, based on a molecular dynamics trajectory. The spatial arrangement of nodes in the graph reflects the frequency of transitions: nodes that are closer together correspond to states between which the ligand transitions more frequently. This visualization provides a high-level map of the binding landscape, revealing both the structural connectivity and the dynamic relationships among ligand-bound states.

**Figure 6.**
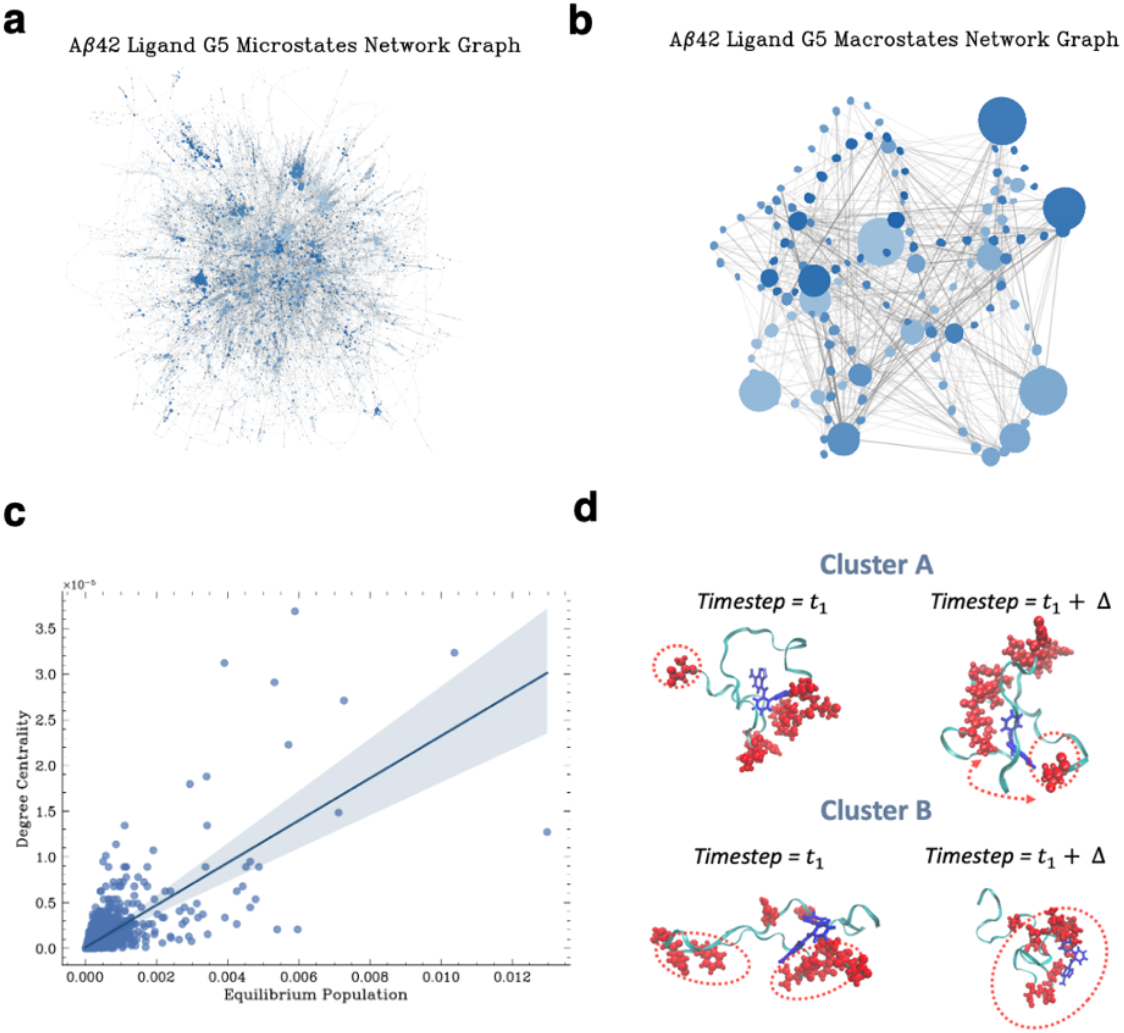
Representation of binding paths on knowledge graphs. **(a, c)** Louvain community detection applied to quintuplet states in Aβ42 (**a)**, with node size proportional to degree centrality. **(b)** Nodes within each community are aggregated into single representative states. Key high-affinity states are identified using dissociation constant (Kd) values. **(c)** Correlation between node centrality and Kd, showing that more highly connected states tend to exhibit stronger ligand binding for Aβ42. **(d)** Representative snapshots from two Aβ42 clusters (Cluster A and Cluster B), highlighting the movement of a flexible arm between the N- and C-termini that facilitates ligand handoff between distant residues.

Quintuplets were clustered using the Louvain algorithm, which is a greedy algorithm that iteratively groups nodes into communities by maximizing modularity, then builds a coarse-grained network of those communities to uncover hierarchical structure^34^. Edges between nodes represent observed transitions between states, and the proximity of nodes reflects the frequency of transitions between them. Nodes were grouped into communities, with each community representing a set of closely connected states (**Figure 6B & Figure 7**). In this way, the original 13,034 states were clustered into 146 macro-states. The size of each node corresponds to the binding affinity (*K*_*D,i*_) of that state (see Eq. 2), providing a visual measure of how strongly each quintuplet interacts with the ligand (**Figure 6C**).

**Figure 7.**
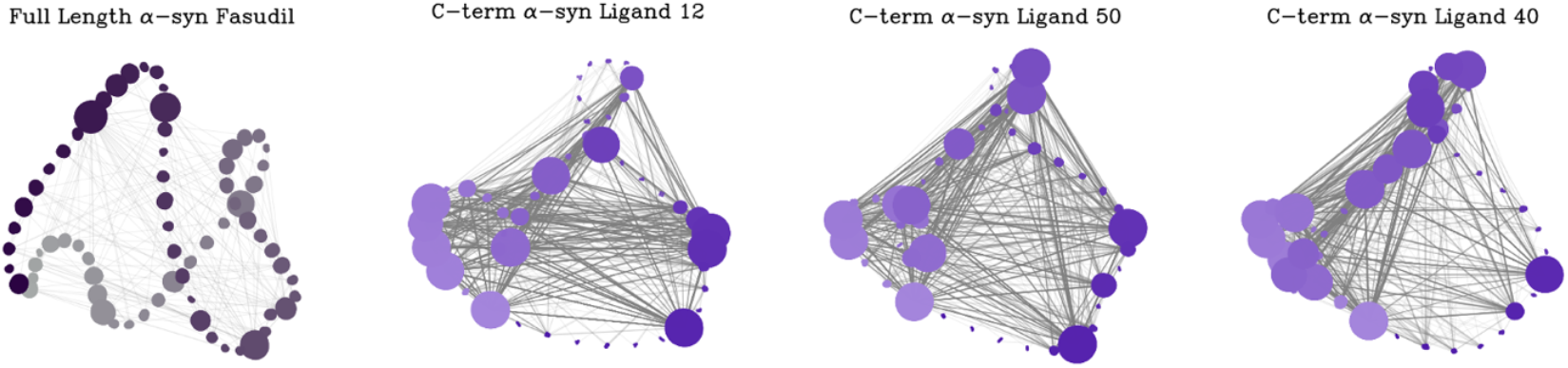
Macrostates network of α-synuclein binding pathways with small-molecule ligands. Binding pathways of the ligands Fasudil, 40, 50, and 12 on full-length α-synuclein (140 residues) and on its C-terminal region, respectively.

This graph-based representation highlights both the dynamics of state transitions and the affinity landscape of the system. Notably, we observed that the closeness centrality of a node, reflecting how centrally positioned a state is within the network, correlates with its binding affinity: states that are more centrally located tend to exhibit stronger ligand binding (**Figure 6C**). Quintuplets were chosen as the representative unit for this analysis based on the outcome of the Chapman-Kolmogorov (CK) test, which indicated that five-residue clusters provided the most robust model.

Additionally, we assessed the intra- and inter-cluster radius of gyration (Rg) and root-mean-square deviation (RMSD) to examine structural variability between and within clusters (**Figure S4**).

### Calculation of the overall dissociation constant from equilibrium populations

Using the equilibrium populations obtained from the stationary distribution of the MSM, we compared the relative dissociation constant (*K*_*D,i*_) for each binding state

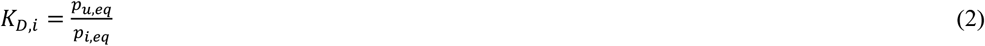

where *p*_*u,eq*_ and *p*_*i,eq*_ are equilibrium probabilities of the unbound state and of state *i*. States with higher equilibrium populations correspond to lower *K*_*D,i*_ values, and therefore stronger binding affinity. For comparison, we calculated the overall binding affinity (Eq. S2), yielding a result of 47 *μ*M, which is consistent with the experimental measurements of 6-40 *μ*M^31,32^.

To assess the robustness of our model across peptides of different lengths, we applied our approach to a previously reported trajectory^28^ of the α-synuclein C-terminus in complex with small molecules. Using the published 60 μs atomistic molecular dynamics simulation performed with the a99SB-disp force field^2^, and following the same analysis protocol described above, we obtained equilibrium dissociation constants of 24.8 µM, 35.8 µM, and 17.6 µM for C-terminal α-synuclein bound to ligands 40, 50, and 12, respectively. To evaluate the dependence of these results on system size, we further applied the same analysis to full-length α-synuclein in complex with the small molecule fasudil, yielding an equilibrium dissociation constant of 59.8 µM **(Figure 7)**.

This analysis enables a quantitative ranking of binding states according to their stability and affinity, providing a thermodynamic interpretation of the interaction of a ligand with the disordered protein. Moreover, it captures the collective behavior of multiple states, including self-transitions (where the ligand remains in the same state) and the possibility of the ligand being bound to multiple residue clusters simultaneously.

## Discussion

In this study, we investigated the complex and dynamic nature of small molecule binding to disordered proteins, focusing on the interaction between Aβ42 and the small molecule G5. Our analysis addresses the challenges in applying to disordered proteins traditional drug discovery approaches, which typically rely on identifying stable binding pockets. These methods are often inadequate for disordered proteins due to their inherent lack of persistent structural features.

Towards this goal, we developed a MSM framework that defines the binding mechanism in terms of a series of transient binding pockets interconnected through a network of transition pathways. Our results show that the formation of a disordered complex between a small molecule to a disordered proteins can be effectively captured using this stochastic approach. In the case of Aβ42, we identified regions on the protein surface where the ligand frequently interacts with specific clusters of residues. Rather than binding to a fixed site, G5 exhibits a diffusion behavior, transitioning smoothly between related residue clusters.

By employing a stochastic modeling approach in terms of an MSM, we quantified both the binding affinities and the transition frequencies between these dynamic states, providing a comprehensive view of the behavior of the ligand. Visualizing the MSM as a knowledge graph further revealed the intricate relationships among binding states and illustrated the overall structure of the binding landscape. The integration of long-timescale molecular dynamics simulations, community detection algorithms, and Markov state modeling underscores the effectiveness of combining advanced computational techniques to study disordered proteins. This approach identifies dynamic binding hotspots and offers a quantitative framework for analyzing both the thermodynamic and kinetic aspects of ligand binding.

The idea of binding paths could help understand binding specificity and suggest how even highly dynamic, disordered proteins can be selectively targeted. Rather than targeting a single, well-defined pocket, a ligand can be designed to follow the characteristic sequence of transient binding pockets that a particular disordered protein presents as it moves. During these motions, the surface of each protein encodes a unique order and spacing of these fleeting contact clusters. By designing a small molecule whose chemical groups engage those clusters in just the right sequence, one can build a filter against off-target interactions. Moreover, the specificity conferred by binding paths derives not only from thermodynamic complementarity but also from kinetic matching. A ligand whose on- and off-rates are tuned to the measured transition frequencies between hotspots becomes effectively bound on its intended path. It also samples the surfaces of other proteins too inefficiently to bind appreciably. Finally, once the high-probability binding paths of a target are computationally mapped, one can extract a fingerprint of residue-transition patterns. This fingerprint then guides fragment-based growth or focused library screening toward chemotypes uniquely matched to that path. In this way, dynamic binding paths become not a barrier to specificity but its very foundation, turning the apparent disorder of the protein surface into a programmable map for highly selective small-molecule recognition.

To provide more context to our findings, we note the analogy between our results and those reported in a recent article that presented a t-SNE-based clustering framework for analyzing the conformational ensembles of disordered proteins and their interactions with small molecules^30^. To further assess the generality of the binding-path framework, we applied the same analysis to α-synuclein systems of different sizes and dynamical regimes, including both the C-terminal region and the full-length protein (Figure 7). A key conclusion of that study was that, despite the structural heterogeneity of disordered proteins, ligand binding can lead to the emergence of metastable substates that are structurally and energetically distinct. In the case of Aβ42, they found that G5 bound to specific, structured substates with characteristic contact patterns and distinct binding energies. In contrast, binding to the C-terminal region of α-synuclein was governed by a dynamic shuttling mechanism, where ligands sample similar interactions across a broad, low-barrier free energy landscape, with moderate correlations between structural features (e.g., bend angle) and binding affinity^30^. Those findings are consistent with our binding path framework. While we emphasize kinetic modeling using MSMs to describe the stochastic evolution of binding contacts, that work focussed on clustering and thermodynamic characterization of conformational ensembles^30^. Both studies converge on the conclusion that ligand binding to disordered proteins involves multiple, kinetically or structurally linked states, and both highlight the relevance of aromatic residues and surface accessibility in determining binding specificity. The results reported in that work further reinforce the notion that binding can be selective without being rigid, supporting the idea that recognition in disordered proteins relies on a continuum of favorable interactions rather than well-defined pockets^30^, an insight foundational to the binding path concept.

We also note another recent study that demonstrated the utility of MSMs to elucidate the folding-upon-binding mechanisms of disordered proteins, specifically in the case of the measles virus NTAIL domain binding to its structured partner XD^35^. Their MSM, constructed using a physically constrained VAMPNet with a multi-input neural network architecture, captures a network of kinetically distinct conformational intermediates characterized by progressive *α*-helix formation and native contact stabilization. In contrast, our MSM framework, designed to study small molecule interactions with disordered proteins such as Aβ42 does not resolve folding events but instead focuses on the stochastic rearrangement of residue contacts, termed binding paths, that describe surface diffusion-like ligand motion. While that MSM is suited to systems involving substantial structural reorganization upon binding, our MSM is optimized for ligands interacting with highly dynamic, unfolded ensembles where binding does not induce folding. Together, these two frameworks underscore the mechanistic diversity of ligand interactions with disordered proteins and highlight the importance of tailoring MSM constructions to the underlying biophysical context.

We anticipate that these findings will have significant implications for drug discovery targeting disordered proteins. By shifting the analysis from binding pockets to binding paths, we open the possibility of identifying targetable regions that may have been overlooked using conventional methods. This change in perspective may ultimately contribute to the development of more effective therapeutic strategies for currently undruggable protein targets.

## Conclusions

We introduced a stochastic framework for understanding and characterizing the interactions between small molecules and disordered proteins. By shifting from a static view of binding pockets to a dynamic view of binding paths, we offer a perspective that captures the transient and flexible nature of these systems. This approach enhances our ability to design small molecules with high affinity, and possibly high specificity, for disordered proteins, thus opening up promising directions for both fundamental research and the development of novel therapeutic strategies.

## Materials and Methods

### Analysis of binding pockets using VAMPNet

Clustering binding pockets using VAMPNet involves a systematic integration of machine learning and dynamic modeling^33^. The input to this process consists of time-series data representing the distances between the center of mass of the ligand and each Cα atom of Aβ42 throughout a 28 µs molecular dynamics simulation of the peptide in complex with the ligand G5. VAMPNet is used to extract slow collective variables (CVs) that capture the essential dynamical features of the system. These CVs are learned by training a neural network specifically optimized to model slow transitions between states. Once identified, the CVs are projected into a latent space, where transitions between configurations are approximated using a Markov process. This enables a kinetic description of the underlying conformational dynamics of the system. Clustering is then performed hierarchically in this latent space. The latent space is first divided into four coarse-grained clusters, which correspond to metastable states of the binding pocket. Each of these primary clusters is further subdivided into finer clusters to capture more granular features of the binding interactions (**Figure 1**). This hierarchical approach facilitates an efficient and dynamic classification of binding pocket configurations and provides insight into their structural and functional diversity. The VAMPNet model was trained in two stages: (i) The first stage used a lag time of 10, 4 states, a learning rate of 5 × 10^-3^, and 30 epochs, with a batch size of 10,000 (**Figure S5**). (ii) The second stage, which refined the model hierarchically, used a lag time of 5, the same number of states and learning rate, and 30 epochs, but with a reduced batch size of 500. These parameters were found to produce the most reliable clustering performance for this system. To assess the reproducibility of the VAMPNet clustering, we performed five independent training runs and compared the resulting state assignments using Adjusted Mutual Information scores (**Figure S6**).

### Calculation of the handoff matrix

Protein-ligand interactions were analyzed using molecular dynamics simulations. Protein structures were processed with the MDAnalysis library to isolate the ligand and the relevant protein residues. To identify contacts, we measured the distances between all atoms of the ligand and protein in each simulation frame. A contact was defined as present when the distance between any two atoms (one from the ligand and one from a residue) was less than 3 Å **(Figure 3)**. To characterize how the ligand moves between different binding regions on the protein surface, we divided the analysis into three distinct cases (**Figure 3**): Case A: Handoff events in which residue *i* was in contact with the ligand at time *t* and residue *j* was in contact at time *t+1*, regardless of whether residue *i* maintained its contact. Case B: Handoff events where residue *i* lost contact with the ligand and residue *j* gained contact at time *t+1*. Case C: Events where residue *i* retained contact with the ligand, while residue *j* additionally gained contact at time *t+1*. These transitions were compiled into handoff matrices, which summarize the frequency of contact transitions between residue pairs. The matrices were visualized using heatmaps, providing insight into the dynamics of ligand movement and the identification of binding hotspots on the disordered protein surface.

### Clustering of residues via enumeration

States in the MSM are determined using a clustering-by-enumeration approach, which involves identifying residue groups (n-plets) that are in contact with the ligand at each time step based on spatial proximity. To carry out this analysis, we processed the protein-ligand interaction data by computing pairwise distances between ligand atoms and protein residues across all frames of a molecular dynamics simulation. A contact was considered present if the distance between any pair of atoms fell below a defined cutoff. Two sources of distance information, one based on the center of mass of the ligand, and the other on proximity to specific residues, were combined using weighted sums to enhance the resolution of contact detection. For each time frame, we then calculated the number of contacts and identified the most frequent n-plet type (i.e., a group of 2, 3, 4, or 5 residues in simultaneous contact with the ligand). These n-plets were considered discrete states, and a transition matrix was built by counting the frequency of transitions between them over time. To correct for enhanced sampling biases in the Aβ42 simulations, transition counts were weighted by the corresponding frame weights (see Supplementary Information). To improve the quality of the model, duplicate states were filtered out, and an explicit unbound state was introduced at both the beginning and end of the trajectory to avoid creating artificially absorbing states. This enumeration-based approach provides a detailed and systematic way to define dynamic contact states, forming the basis for analyzing state transitions and quantifying binding affinities in disordered protein-ligand systems.

### Solving the master equation

In this analysis, we describe a MSM that represents the transitions between different binding states, each defined as an n-plet of residues (e.g., quadruplets or quintuplets) simultaneously interacting with the ligand. Each binding state is assigned an index *i* = 1,…,n, and the probability of the system being in state *i* at time *t* is denoted as *p*_*i*_(t). These probabilities are assembled into a state probability vector P(t) = [*p*_*i*_(t), …, *p*_*n*_(t)]. The time evolution of this probability vector is governed by the master equation (Eq. 1), which is a system of linear differential equations that describes how the population of each binding state changes over time due to stochastic transitions. We assume that the dynamics are Markovian (i.e., memoryless), time-independent, and linear, with constant transition rates. Under these conditions, the system evolves towards a stationary distribution *Peq*, which corresponds to the eigenvector associated with the zero eigenvalue of the generator matrix L. Each component *p*_*i*_*eq* represents the equilibrium probability of the system being in state *i*. To estimate the transition rates from simulation data, we evaluate how the population vector *P*(*t*) changes after a small time increment Δ*t*. This yields an approximate discrete-time formulation:

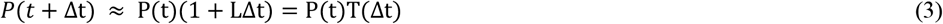

Here, T(Δt) is the transition probability matrix, and its dominant eigenvalue (equal to 1) corresponds to the equilibrium distribution.

### Graph network clustering

Based on the results of the Chapman-Kolmogorov test, which indicated that models using quintuplets provided the most stable MSM, we constructed a network graph G to visualize the connectivity between binding states. This graph was created using the NetworkX library, a Python library for creating, manipulating, and analyzing complex networks (graphs), providing data structures for nodes and edges as well as a wide range of built-in algorithms for tasks like path finding, clustering, and network statistics. In our graph, each node representing a quintuplet state, and each weighted edge representing the transition frequency between two states (**Figure 6a**). To prevent artifacts, self-loops (transitions from a state to itself) were removed from the graph. We applied Louvain clustering (with resolution = 3) to identify communities of closely connected states. Each node was colored according to its assigned community. The size of each node was scaled by its degree centrality, which was previously shown to correlate with the state’s binding affinity (**Figure 6c**). To enhance visual clarity, node positions were arranged using a spring layout, and edges were drawn with reduced opacity to avoid clutter. To quantify the importance of each community, we summed the equilibrium probabilities Peq of all nodes within that community. A sigmoid function was then applied to these summed probabilities to scale the visual representation of community importance (see Supplementary Information). Finally, a supergraph was constructed to connect the communities, with individual subgraphs positioned relative to the supergraph layout (**Figure 6b & Figure 7**), providing a high-level view of how groups of binding states are organized and interrelated in the MSM.

## Data Availability

https://github.com/louetadelie1/MSM.git

## Acknowledgements

We would like to acknowledge funding from UKRI (10059436 and 10061100, M.V.).

## Author contributions

A.L., G.H. and M.V. conceived the project. A.L. performed the computational work. G.H. and M.V. funded the project. A.L., G.H. and M.V. analyzed data and wrote the article.

## Competing interests

The authors declare no competing interests.

## Notes

https://github.com/louetadelie1/MSM.git

### Competing Interest Statement

The authors have declared no competing interest.

### Summary of Updates

We have added more data to the original manuscript.

