## Supplemental Information for "Binding Paths: Describing Small Molecule Interactions with Disordered Proteins via a Markov State Model"

**Aβ42 simulations**

All-atom metadynamic metainference simulations of free and bound Aβ42 were carried out using GROMACS 2018.3 with PLUMED 2.6.0-dev, employing the CHARMM22* force field and TIP3P water model, following a previously reported protocol[1]. Forty-eight initial conformations were generated and equilibrated at 278 K, first for 500 ps in the NVT ensemble using the Bussi thermostat, then for 500 ps in the NPT ensemble using Berendsen pressure coupling[1]. Production runs were conducted in the NPT ensemble with the Parrinello-Rahman barostat[1]. Enhanced sampling was achieved via parallel bias metadynamics using well-tempered and multiple-walker schemes[1]. Bias factors were set to 24 and 49 for unbound and bound simulations, respectively[1]. The unbound system used six collective variables (CVs) for conformational sampling, while the bound system included 14 additional CVs for protein-ligand contacts and four for ligand dihedrals. Simulations totalled 27.8 μs (unbound) and 28.2 μs (bound).

**α-Synuclein simulations**

To assess the robustness of our model across peptides of different lengths, we applied our approach to a previously reported trajectory[2] of the α-synuclein C-terminus in complex with small molecules. Using the published 60 μs atomistic molecular dynamics simulation performed with the a99SB-disp force field[2], and following the same analysis protocol described above, we identified 38, 37, 39, 68 residue clusters and reconstructed the binding paths of three arbitrarily selected small molecules (ligands 40, 50, 12, and Fasudil) across the α-synuclein surface.

**Sigmoid function for data transformation**

To determine the most suitable sigmoid transformation for visualizing the relative importance of different communities in our dataset, we first constructed an inverse mapping of state groupings, clustering related elements together. For each group, the equilibrium probabilities (Peq) of its constituent states were summed to generate input values for the transformation. We applied a sigmoid function of the form:

$$S(x) = \frac{A}{1+\exp(-k(x-x0))} \qquad \text{(S1)}$$

where A is the amplitude (scaling factor), k controls the steepness of the curve, and $x_0$ is the midpoint of the transition. Several parameter sets were tested to evaluate their effect on the transformed values. These outputs were visualized using scatter plots to compare the behavior of each transformation function and assess their ability to reflect meaningful differences in community importance (Figure S1). The best-fitting parameters for our data were determined to be A = 1000, x0 = 200, and k = 0.025. This parameter set provided the most effective scaling, clearly differentiating communities while preserving the overall distribution characteristics of the original equilibrium probabilities.

**Binding affinity calculations**

The binding affinity was estimated from molecular dynamics (MD) simulations using two approaches: a frame-based occupancy method and a kinetics-based method. Both methods were applied across the full system without assuming predefined binding pocket locations. The dissociation constant $K_D$ was estimated by comparing the number of frames where the ligand was unbound ($P_u$) versus bound ($P_b$), and scaling by the simulation box volume $V$

$$K_D = \left(\frac{P_u}{P_b}\right)\frac{1}{N_A \cdot V} \tag{S2}$$

where $N_A$ is the Avogadro number.

For comparison, we then estimated $K_D$ based on the average residence times the ligand remained in bound and unbound states (**Table S1**). The alternative estimation was based on the average residence times the ligand remained in bound and unbound states. Consecutive frames with non-zero contacts were treated as bound intervals ($\tau_b$), while those with zero contacts were unbound ($\tau_u$)

$$K_D = \left(\frac{\tau_u}{\tau_b}\right)\frac{1}{N_A \cdot V} \tag{S3}$$

**VAMPNet validation**

The first VAMPNet run was performed with a lag time of 5 steps, targeting a reduced model of 4 macrostates. The network was trained for 30 epochs using a learning rate of $5 \times 10^{-3}$ and a batch size of 10,000. A second VAMPNet refinement was then conducted hierarchically using the output of the first model. This second run used a lag time of 10 steps with the same number of target states (4), learning rate ($5 \times 10^{-3}$), and number of epochs (30), but employed a smaller batch size of 1,000 to better capture finer features in the state transitions. Cluster output from the second round of VAMPNet training (lag time = 10) after 5 independent runs **(Figure S5)**. The plot shows the consistency of state assignments within this cluster across runs. Adjusted mutual information (AMI) scores were computed to quantify agreement between runs, indicating stable and reproducible clustering behavior.

**Reweighting simulations**

To reweight the biased trajectories, the time-ordered trajectories from each replica were first combined into a single continuous trajectory using the gmx trjcat tool. Parallel-bias metadynamics (PBMetaD) was used to apply biases, with the corresponding HILLS files capturing the Gaussians deposited by each replica at every simulation step. Reweighting was then carried out on the full concatenated trajectory using plumed driver, resulting in a weight assigned to each frame. This method ensures alignment between the trajectory of each replica and its bias history, enabling accurate computation of unbiased weights.

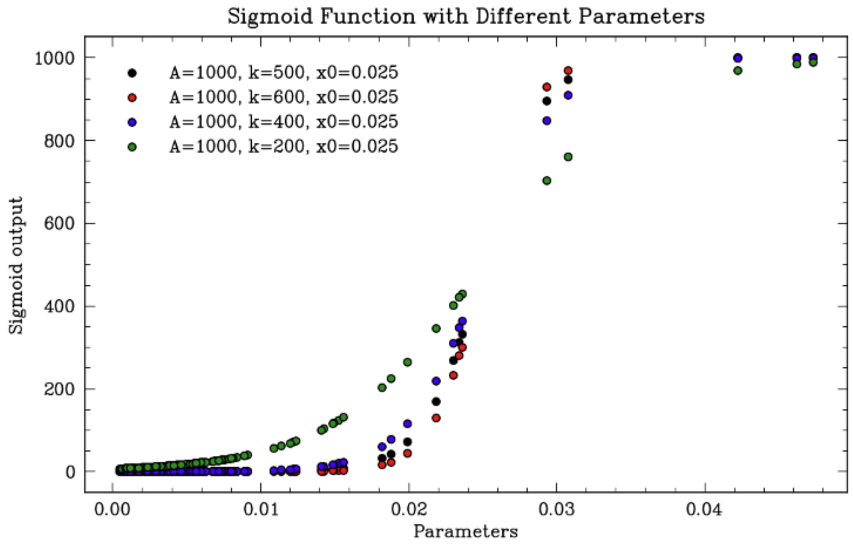


**Figure S1. Testing Sigmoid Parameters for data transformation.**

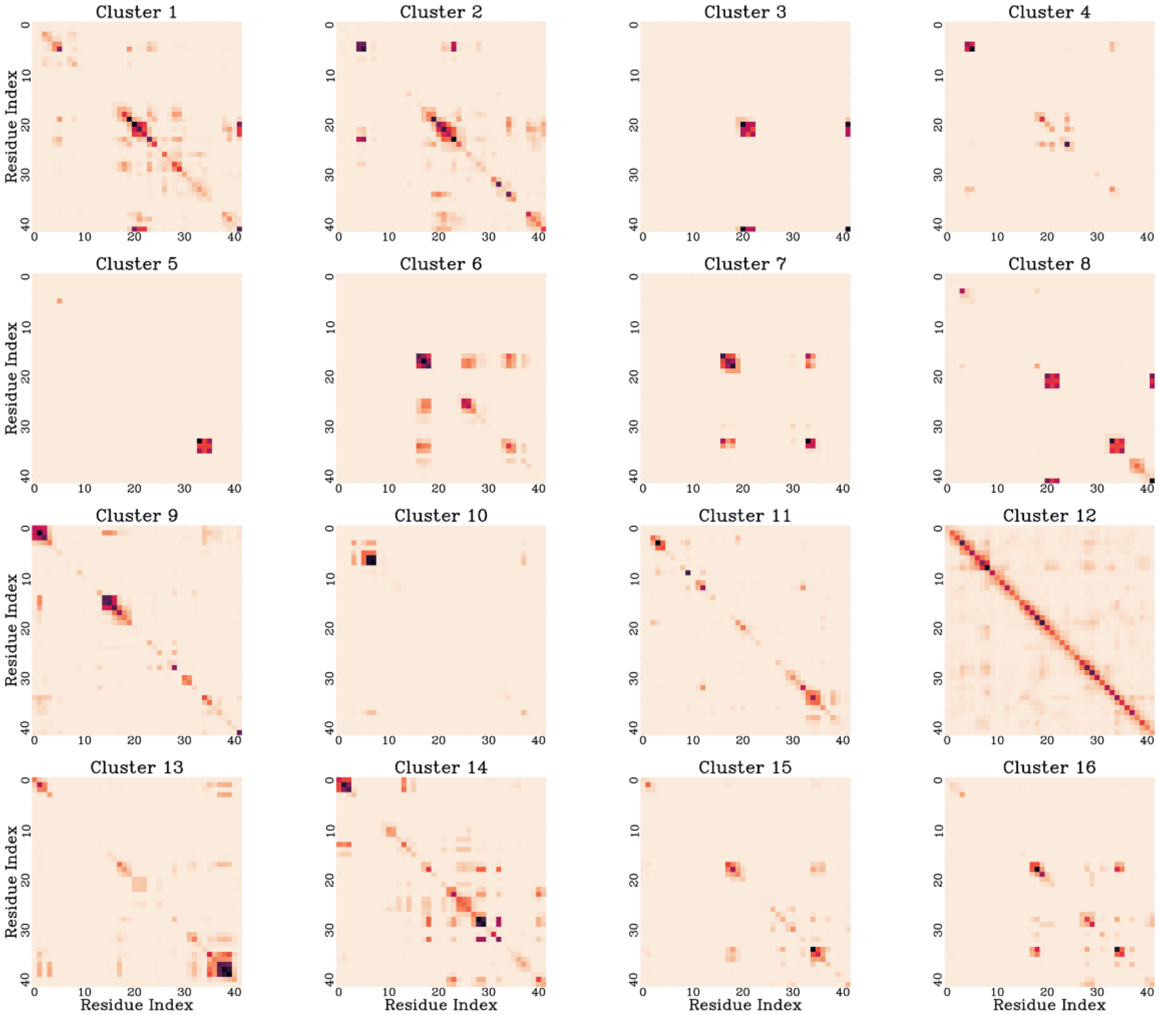


**Figure S2**. **Population distribution of the 16 microstates obtained from the initial VAMPNet clustering of the Aβ42-G5 complex.** The majority of frames are assigned to state 16, indicating the dominance of a single contact configuration and highlighting the limitations of conventional clustering methods for this highly dynamical, disordered system.

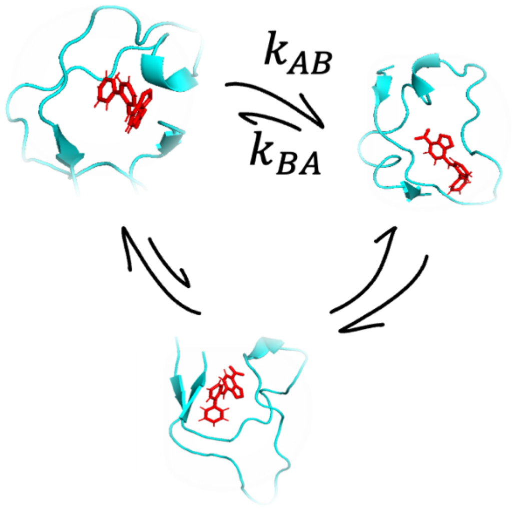


**Figure S3:** Illustration of the Markov state model.

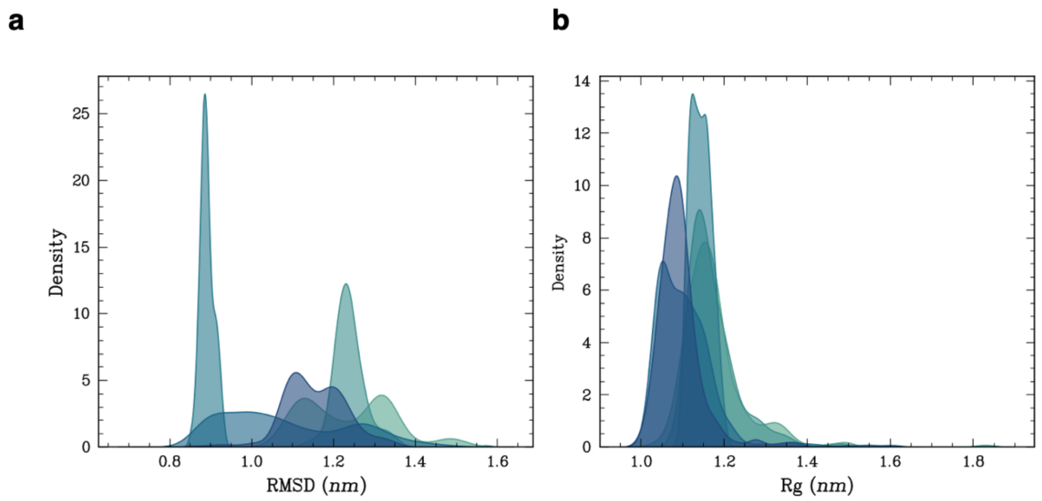


**Figure S4. Stochastic steps along the binding paths.** (**a,b**) Histograms showing the probability density distributions of the radius of gyration (Rg) (a) and root-mean-square deviation (RMSD) (b) for five representative clusters. The results illustrate intra- and inter-cluster variability in the system. The RMSD inter-cluster mean difference is 0.160 nm and the intra-cluster standard deviation averages 0.130 nm; the Rg inter-cluster difference is 0.110 nm and the intra-cluster standard deviation is 0.111 nm.

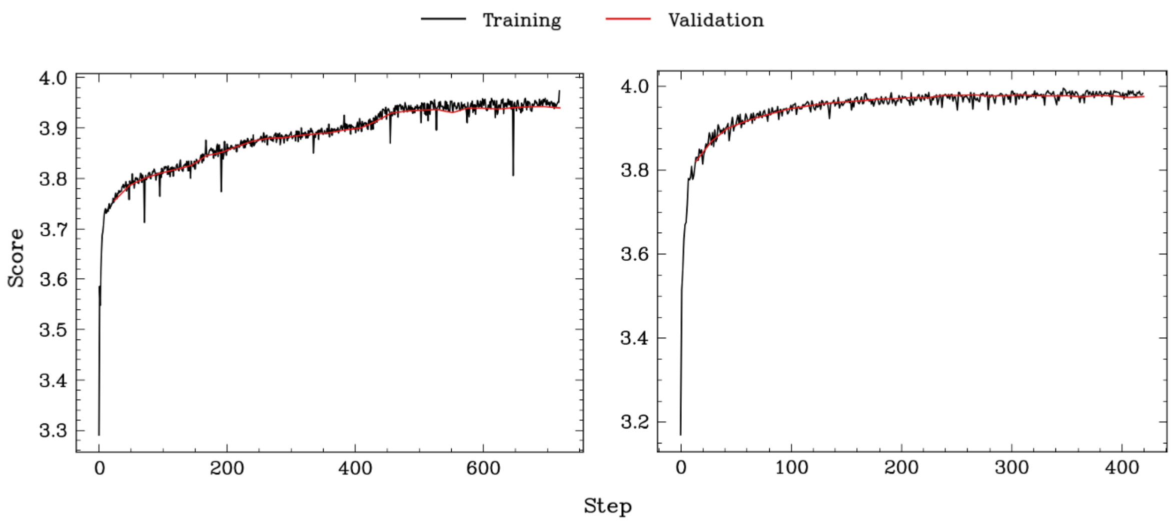


**Figure S5. Training and validation scores during VAMPNet optimization (**for 1 run**)**, plotted on a log-log scale. The VAMP2 score is shown as a function of training step for both training and validation datasets, indicating convergence behavior and model performance. This was assessed over both iterations in the hierarchical VAMPNET scheme. The first plot **(left)** shows the training/validation for the first hierarchical run, and the second plot **(right)** shows the training/validation for the second hierarchical run.

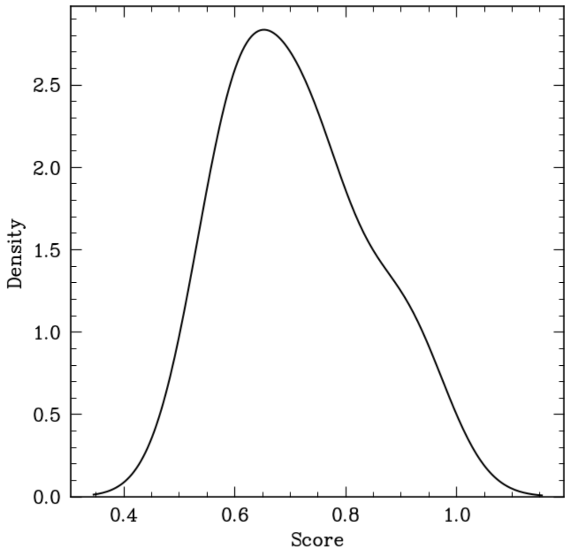


**Figure S6. Reproducibility of VAMPNet clustering across five independent training runs.** The plot shows the adjusted mutual information (AMI) scores between clustering outcomes, quantifying the consistency of state assignments. High AMI values indicate stable and reliable identification of metastable states in the Aβ42-G5 system.

**Table S1. Transition labels between state numbers.** This table lists the top transitions between quintuplet states, showing both state identifiers and the corresponding residue groupings involved in each transition

| Transition Label | From State # | From State Quintuplet | To State # | To State Quintuplet |
|---|---|---|---|---|
| T1 | 5 | (4, 6, 7, 8, 30) | 5 | (4, 6, 7, 8, 30) |
| T2 | 11 | (6, 7, 8, 39, 40) | 11 | (6, 7, 8, 39, 40) |
| T3 | 28 | (30, 31, 34, 35, 41) | 28 | (30, 31, 34, 35, 41) |
| T4 | 3 | (3, 7, 11, 14, 37) | 3 | (3, 7, 11, 14, 37) |
| T5 | 1 | (2, 3, 23, 36, 37) | 1 | (2, 3, 23, 36, 37) |
| T6 | 21 | (9, 15, 16, 34, 39) | 21 | (9, 15, 16, 34, 39) |
| T7 | 4 | (3, 36, 37, 38, 40) | 4 | (3, 36, 37, 38, 40) |
| T8 | 14 | (7, 11, 14, 31, 37) | 14 | (7, 11, 14, 31, 37) |
| T9 | 17 | (8, 12, 29, 30, 33) | 17 | (8, 12, 29, 30, 33) |
| T10 | 22 | (9, 15, 16, 34, 40) | 22 | (9, 15, 16, 34, 40) |
| T11 | 6 | (4, 6, 7, 8, 40) | 5 | (4, 6, 7, 8, 30) |
| T12 | 5 | (4, 6, 7, 8, 30) | 6 | (4, 6, 7, 8, 40) |
| T13 | 14 | (7, 11, 14, 31, 37) | 3 | (3, 7, 11, 14, 37) |
| T14 | 3 | (3, 7, 11, 14, 37) | 14 | (7, 11, 14, 31, 37) |
| T15 | 7 | (4, 17, 20, 22, 41) | 8 | (4, 17, 20, 38, 41) |
| T16 | 8 | (4, 17, 20, 38, 41) | 7 | (4, 17, 20, 22, 41) |
| T17 | 24 | (9, 19, 21, 22, 30) | 27 | (19, 20, 21, 22, 30) |
| T18 | 22 | (9, 15, 16, 34, 40) | 20 | (9, 13, 15, 16, 34) |
| T19 | 18 | (8, 18, 29, 30, 33) | 17 | (8, 12, 29, 30, 33) |
| T20 | 27 | (19, 20, 21, 22, 30) | 24 | (9, 19, 21, 22, 30) |
| T21 | 9 | (4, 19, 20, 22, 41) | 8 | (4, 17, 20, 38, 41) |
| T22 | 23 | (9, 19, 21, 22, 27) | 27 | (19, 20, 21, 22, 30) |
| T23 | 16 | (8, 9, 13, 16, 19) | 19 | (9, 12, 13, 16, 17) |
| T24 | 8 | (4, 17, 20, 38, 41) | 9 | (4, 19, 20, 22, 41) |
| T25 | 15 | (8, 9, 12, 29, 33) | 18 | (8, 18, 29, 30, 33) |
| T26 | 2 | (3, 6, 7, 11, 14) | 14 | (7, 11, 14, 31, 37) |
| T27 | 14 | (7, 11, 14, 31, 37) | 2 | (3, 6, 7, 11, 14) |
| T28 | 13 | (7, 8, 12, 28, 35) | 12 | (7, 8, 10, 12, 25) |
| T29 | 26 | (16, 17, 24, 31, 34) | 25 | (16, 17, 20, 23, 24) |
| T30 | 10 | (4, 19, 20, 38, 41) | 7 | (4, 17, 20, 22, 41) |
